# Impacts of water quality on *Acropora* coral settlement: The relative importance of substrate quality and light

**DOI:** 10.1101/2020.11.19.390724

**Authors:** Gerard F. Ricardo, Charlotte E. Harper, Andrew P. Negri, Heidi M. Luter, Muhammad Azmi Abdul Wahab, Ross J. Jones

**Affiliations:** Australian Institute of Marine Science, Townsville, 4810, Queensland, Perth, 6009, Western Australia, and Darwin, 0810, Northern Territory, Australia; University of Plymouth, Drake Circus, Plymouth, Devon, PL4 8AA, UK

**Keywords:** Coral settlement, water quality, sediment, spectral patterns, light intensity

## Abstract

Coral larval settlement patterns are influenced by a vast array of factors; however, the relative roles of individual factors are rarely tested in isolation, leading to confusion about which are most crucial for settlement. For example, direct effects of light environment are often cited as a major determinate of settlement patterns, yet this has not been demonstrated under environmentally realistic lighting regimes in the absence of confounding factors. Here we apply programmable multispectral lights to create realistic light spectra, while removing correlating (but not obvious) factors that are common in laboratory settlement experiments. Using two common species of *Acropora* – key framework builders of the Great Barrier Reef – we find little evidence that light intensity or changes in the spectral profile play a substantial role in larval settlement under most environmentally realistic settings but can under more extreme or artificial settings. We alternatively hypothesise and provide evidence that chronic conditions of light and recent sediment exposures that impact benthic substrates (e.g., crustose coralline algae) preceding settlement have a greater impact, with up to 74% decrease in settlement observed on substrates with prior exposure and poor water quality conditions. Management of water quality conditions that impact the quality of benthic-settlement substrates therefore should present a priority area of focus for improving coral recruitment.

## Introduction

Understanding the relative importance of drivers affecting recruitment has been a fundamental objective of marine ecology for many decades (Giese 1959, Roughgarden et al. 1988), and is of critical importance for predicting how reefs will recover following disturbances, and aiding intervention efforts such as translocation of progeny to damaged reefs (Randall et al. 2020, Vanderklift et al. 2020). In coral reef ecosystems, direct light and spectral quality are perceived as key influences on coral settlement patterns, with implications for depth-zonation in corals as well as changes in settlement preferences under poor water quality regimes (Mundy & Babcock 1998, Mundy & Babcock 2000, Fox 2004, Strader et al. 2015, Turner et al. 2018). In the shallows, the water column and benthos are exposed to high irradiance across a broad spectrum; but the spectral profile shifts left across the visible spectrum with increasing depth to become dominated by shorter (blue) wavelengths (Kirk 1994). Across sediment gradients, there is a competing right-shift towards the centre of the visible spectrum with green-yellow light becoming more dominant (Jones et al. 2016, Strydom et al. 2017, Cussioli et al. 2020). Under both gradients, these shifts in the spectral profile co-occur with decreasing light intensity.

For adult corals, there is a strong mechanistic basis for changes in the light environment influencing individual health. Adult corals are primarily autotrophic during their benthic lifecycle stage (Muscatine 1990), and coral photosynthesis and growth may be limited by decreases in light intensity and spectral shifts away from the maximally absorbance of regions of chlorophyll *a* (blue and red wavelengths) termed ‘photosynthetically usable radiation’ (PUR) (Morel 1978, Dustan 1982). Consequently, benthic-light monitoring is suggested as a valuable metric to measure and manage water quality for inshore reefs (Gruber et al. 2019, Magno-Canto et al. 2019, Robson et al. 2019, Jones et al. 2020). But for coral early-life stages preceding the establishment of autotrophy (i.e., acquisition and utilisation of algal symbionts as the primary energetic pathway), the contribution of direct light influencing recruitment patterns is an area of active research. Corals settle non-randomly, using a range of cues which is thought to increase their chances of survival through the post-settlement stage when mortality is high (Doropoulos et al. 2016, Brunner et al. 2020). Numerous studies have suggested light quality and quantity affects where larvae settle at depth and at which orientation (Mundy & Babcock 1998, Mundy & Babcock 2000). Indeed, sensory mechanisms to detect changes in light quality and intensity have been proposed with evidence of opsins and cryptochromes (a class of proteins involved in circadian rhythms) (Levy et al. 2007, Mason et al. 2012).

Despite plausible mechanisms to sense light in larvae, settlement responses to light often contradict, even within the same species. For example, *Acropora tenuis* settlement patterns purportedly increase (Mundy & Babcock 1998), decrease (Suzuki & Hayashibara 2006), or remain relatively unchanged (Yusuf et al. 2019) with light intensity. However, an issue is that many factors within the water column and on the benthos may simultaneously act on a settling larva, and that these are rarely controlled in experiments making interpretation of settlement patterns difficult. For example, grooves, edges, and surface orientations of the substrate all influence the micro-light environment relevant to a larva. However, these physical characteristics can also affect thigmotactic behaviour in larvae, potentially creating unintended artefacts in settlement studies attempting to assess the influence of light.

Therefore, even though light patterns at the time of settlement may correlate with settlement behaviour, it may not be a direct driver of observed patterns, as suggested for temperate invertebrate communities (Irving & Connell 2002). Further, light treatments common in laboratory experiments often do not match those that occur *in situ*. Most use narrow-band (monochromatic) lighting or aquarium lights that exploit PUR by matching the profile to the absorption spectrum of the symbionts (Jones et al. 2020). While these systems are simple or optimized for photosynthesis and growth, the conditions do not effectively mimic the broader spectral patterns that are prevalent in the water column, nor the spectra that result from shifts in deeper water and in the presence of suspended sediments or plankton.

An alternate, but often overlooked, explanation of settlement patterns across depths and orientations is the potential influence of environmental conditions on the benthic substrate community in the weeks to months prior to settlement (Baird et al. 2003). Often the substrate in field recruitment tile surveys and laboratory experiments are conditioned in the same environments and configurations as those exposed to the larvae during settlement, making it challenging to disentangle long-term environmental and water-quality factors acting on the substrate from the acute light environment. For example, upwards-facing surfaces are exposed to sedimentation and higher light which in turn structures the substrate community. Autotrophs are common on these surfaces, whereas low-light specialists and heterotrophs often dominate undersides (Erwin et al. 2008, Doropoulos et al. 2020). Therefore, biotic factors that attract and repel larvae on these surfaces may impact settlement to a greater degree than the direct light environment present at the time of settlement. Conversely, field studies that attempt to link the substrate community with settlement patterns often ignore the water-quality conditions at the time of settlement, and can suffer from the known issues with identifying settlement cues and inhibitors visually (Erwin et al. 2008).

Recently, technological advancements in *in situ* spectroradiometers and aquarium light emitting diode (LED) technology have allowed for improved characterisation and simulation of a range of natural water column spectra and intensities. Here, we employ these technologies to address some of the limitations of prior light experiments by applying highly controlled but realistic experimental approaches to investigate environmental influences on settlement. We first measure *in situ* exposures with spectroradiometers at scales relevant to settling larvae across depth, turbidity and within crevices. Next, we use these observations to inform the simulation of a range of natural water column spectra and intensities using a custom-built multispectral feedback LED lighting system for application in coral settlement experiments. These experiments are designed to eliminate variables such as three-dimensional shading and edge-effects that have confounded most previous experiments. We hypothesise that depth and water-quality light profiles at the time of settlement will lead to changes in settlement success. As an alternative hypothesis, we assess whether chronic (longer-term) water-quality conditions that impact the conditioning and quality (attraction or inhibition) of the settlement substrate is the primary driver of settlement success.

## Methods

### Methodological overview

To investigate the influence of light (quality and quantity) on coral settlement, coral larvae from the common Indo-Pacific corals *Acropora millepora* and *Acropora tenuis* were exposed to several different light treatments in a series of experiments. Light treatments included different intensity and spectra of narrow-band monochromatic light as well as spectra corresponding to clearer-water and naturally turbid coastal environments (see Settlement series 1; Table 1). Next, substrates conditioned at two sites of differing water quality were used to assess the impact of chronic effects of exposures on the substrate and the subsequent impact on coral settlement (see Settlement Series 2; Table 1).

**Table 1.** Experimental treatments used in each experiment. PAR = Photosynthetically Active Radiation (μmol photons m^−2^ s^−1^). DLI = Daily Light Integral (mol photons m^−2^ d^−1^). ‘Upwards’ and ‘downwards’ refer to orientation of the settlement discs during field conditioning. SS = suspended sediment. NTU = Nephelometric Turbidity Units.

| Design | Exp. | Spectra | PAR | DLI |
| --- | --- | --- | --- | --- |
| *Settlement series 1* |  |  |  |  |
| Monochromatic | Exp.1 & 5 | Broad, blue, green, yellow, red | 30 | 1.3 |
| Simulated depth | Exp.2 | Low broad, high broad, low blue, high blue | 30, 320 | 1.3, 14 |
| Simulated turbidity | Exp.3 and 5 | ~0.8, 1.5, 3, 9 NTU | 30 | 1.3 |
| Light intensity | Exp.4 | Broad and SS-simulated (0.8 and ~9 NTU) | <1, 10, 30, 100, 300 | <0.1, 0.3, 0.9, 3, 9 |
| Experiment |  | Treatment | PAR | DLI |
| *Settlement series 2* |  |  |  |  |
| Deposited sediment | Exp. 6 | 0.1, 0.3, 0.8, 2.5, 8, 25, 83, 250 $\text{mg cm}^{-2}$ | 30 | 1.3 |
| Field conditioning | Exp. 7 | Low-turbidity upwards, low-turbidity downwards, high-turbidity upwards, low-turbidity downwards | 30 | 1.3 |

### Coral collection, larval culturing, and substrate conditioning

Coral collection and larval culturing followed Heyward and Negri (1999) and Ricardo et al. (2018), and further details can be found in the Supplementary Material (Text S1). Artificial substrates for settlement were created using 7-cm diameter polyvinyl chloride (PVC) discs. PVC was used for the following reasons i) it presents a flat surfaces to ensure there is no shading of downwelling light, ii) it can be manufactured to minimise edge-effects, and iii) is successful at conditioning a number of settlement cues and thus frequently used in settlement experiments (Lee et al. 2009, Fabricius et al. 2017, Kennedy et al. 2017, Ricardo et al. 2017). For laboratory experiments, they were conditioned under layers of 70% block shade cloth (maximum Photosynthetically Active Radiation (PAR) light intensity = ~250 μmol photons m^−2^ s^−1^) in flow-through outdoor SeaSim aquaria for ~3 months of the presence of crustose coralline algae (CCA) *Porolithon onkodes* a common species known to induce settlement of *Acropora* coral larvae (Heyward & Negri 1999). The discs were conditioned randomly within the aquaria and CCA covered about 20–40% of the total disc area at the commencement of each experiment.

Before settlement experiments commenced, larvae were assessed for their settlement competency (*sensu* Guest et al. (2010)), and further details can be found in the Supplementary Material (Text S1). For all settlement experiments excluding Exp. 6, each disc was positioned at the base of a 500 mL black chamber containing 250 mL 0.5 μm-filtered seawater (FSW). The outside of the disc was wrapped in thread seal Teflon tape which effectively prevented larvae settling on the conditioned vertical edges of the disc. Additionally, cleaned river sand was added to the bottom of the chambers up to prevent larvae settling in the edges (Ricardo et al. 2017). At the commencement of the experiments, larvae (n = 30) were added into the chambers to settle. A constant light intensity, as opposed to a ramping light intensity, was used to prevent confounding across the light treatments, and each treatment was programmed on a 12:12 light/dark cycle with the larvae censused after 12 hours of light exposure and then after a further 12 hours of darkness to assess any latency effects. All experiments were conducted at 27°C and measured using a Hobo Onset Temperature logger in a reference chamber under each light.

### In situ measurements for laboratory light treatments

*In situ* light measurements of this study were conducted at inshore sites within Cleveland Bay, Great Barrier Reef using a range of PAR light meters and spectroradiometers, with these data used to justify the light treatment-levels used in experiments. Briefly, a light meter connected to a diving pulse amplitude modulating fluorometer (PAM) (Walz, Germany) was used to characterise the benthic substrate light intensity levels that coral larvae are likely to settle (Table 2). We consequently selected 30 μmol photons m^−2^ s^−1^ for the ‘control’ light intensity in experiments. To characterise long-term underwater light intensities and spectral changes with turbidity, two eight-wavelength multispectral radiometers (MS8, In Situ Marine Optics, Australia) were deployed in ~5-m depth at Florence Bay (lower turbidity; 19.121°S, 146.882°E) and Picnic Bay (higher turbidity; 19.186°S, 146.840°E) reefs. Additionally, to characterise more extreme light spectral profiles under turbidity and depth, instantaneous spectral profiles were measured using a USSIMO hyperspectral radiometer (In Situ Marine Optics, Australia) at a clear-water site (19.085°S, 146.947°E) at two depths (5 m and 21 m), and a highly turbid-water site at 5 m depth (19.169°S, 146.905°E). The light spectral profile at 5-m depth (~0.8 NTU) was then applied as the experimental control in all experiments. In addition to the *in situ* spectral measurements described above, individual light channels of the lighting systems (see below) were used to replicate more artificial monochromatic wavelengths commonly applied in spectral-light response studies. Further details can be found in the Supplementary Material (Text S2).

**Table 2.** In situ measurements for design of the laboratory experiments. See text for further site details. PAR = Photosynthetically Active Radiation (μmol photons m^−2^ s^−1^). DLI = Daily Light Integral (mol photons m^−2^ d^−1^). FWHM = Full-width half-maximum.

| Measurement | Instrument | Measurement | Details |
| --- | --- | --- | --- |
| Substrate light intensities | Diving PAM | 15.56 PAR; ~0.5 DLI | Mean discrete measurements over a substrate during midday on a cloudless day. |
| Temporal light intensities | MS8 multispectral | 342.89 midday PAR; 7.8 DLI | Mean intensities over a 3-mo. period at the low-turbidity site. |
| Spectral profiles | USSIMO hyperspectral | 484 nm; 187 FWHM, 0.8 NTU (clear shallow)<br>484 nm; 126 FWHM, 1 NTU (clear deep)<br>559 nm; 124 FWHM, 5 NTU (turbid shallow) | Modal and full-width half maximum wavelength at clear shallow and deep water, and a shallow turbid-water site |
| Turbidity (NTU) | DL3 nephelometer | 0.8 NTU, 1.5 NTU, 3.0 NTU, 9.4 NTU | NTU corresponding to the spectral profiles used. These values represent the 6<sup>th</sup>, 45<sup>th</sup>, 77<sup>th</sup>, and 88<sup>th</sup> percentile recorded at the turbid water site (Picnic Bay). NTU can be converted to suspended sediments using a conversion factor of 1.1. |

\*assumes a cosine approximation

### Experimental lighting setup

Ten spectrally-tuneable LED-based light systems were designed in-house to simulate realistic light spectra found in marine environments. Each of the lights contained 840 individual LEDs controlled by 19 independent colour channels. The lighting system was integrated into a PLC system (Siemens PCS7, Supervisory Control and Data Acquisition System), with the channels adjusted through a feedback system to a user-defined light intensity level. Neutral density shade film was also added to each light to allow for greater control of the channels at lower light intensities. Light intensity was measured with a PAR Quantum sensor (Skye SKP 215). For Daily Light Integral calculations (DLIs), PAR measurements across a 24-hr period were summed. Irradiance in experiments were measured with a Jaz spectrometer (Ocean Optics) calibrated to a recently calibrated HL-2000 lamp. The spectrometer was connected to a 5-m long 400-μm diameter fibre optic with a 3.9 mm diameter in-water cosine corrector attached. Irradiance from spectrometers were interpreted as relative rather than absolute values because of the common issues associated with spectrometer calibrations, light flickering, and attenuation of light under cosine correctors (Johnsen 2012), and were therefore corrected against the PAR Quantum sensor. All absolute irradiance readings (in μW cm^−2^ nm^−1^) were binned to the nearest nanometre and converted to photons (μmol photons m^−2^ s^−1^ nm^−1^) using the equation (Thimijan & Heins 1983):

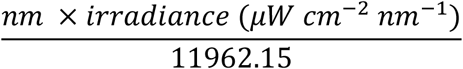

### Settlement Series 1: Acute light exposure–experimental settlement trials

#### Exp. 1. Artificial light spectra on A. millepora settlement

Five replicate chambers were positioned under each light treatment which included a realistic broad-spectrum control and four narrow monochromatic bands. The width of the bands is described using full-width at half maximum (FWHM). The median and FWHM correspond to broad-spectrum: median λ = 535 nm; FWHM = ~172 nm; artificial blue light: λ = 441 nm; ~17 nm, artificial green light: 499 nm; ~25 nm, artificial yellow light: 602 nm; ~28 nm, and artificial red light: 658 nm; ~20 nm) (Table 1). To demonstrate repeatability and to increase sensitivity to treatments, we repeated the experiment using a ‘choice’ design (more details in Supplementary Material, Text S3).

#### Exp. 2: Realistic depth-dependant light on A. millepora settlement

Settlement preferences in response to depth-derived light spectra and intensity is often cited in the literature to influence the zonation of corals (Mundy & Babcock 1998). Broad and artificial blue spectra were supplied at two light intensities (Table 1). The broad-light intensities used were based upon the USSIMO field data described above at 5-m depth and 21-m depth from the clear blue-water site and broadly matched spectral profiles used in Mundy and Babcock (1998). The light channels for the ‘blue’ treatment were used to reproduce more extreme artificial peaks.

#### Exp. 3: Realistic turbidity-dependant light spectra on A. millepora settlement

Five turbidity-dependent light spectral profiles were simulated using NTU percentiles in Jones et al. (2020), with light on a 12:12 h light/dark cycle and PAR held constant (~30 μmol photons m^−2^ s^−1^; 1.3 DLI). The spectral patterns treatments were bracketed between NTU recorded at the clear-water site (site 47, ~0.8 NTU), and the 88^th^ percentile at the turbid-water site (Picnic Bay, 9.4 NTU), which represented the best and worse-case turbidity scenarios (Table 1 & 2). All spectral profiles used were at environmentally realistic light-intensity combinations.

#### Exp. 4: Light intensity in clear and turbid spectra on A. millepora settlement

This experiment was designed to examine the combined effects of changes in light intensity (5 levels) and quality (broad and shifted spectrums that would be associated with suspended sediment concentrations equivalent to 0.8 NTU and ~9 NTU at 5-m depth). Five light intensities were selected to cover the range of environmentally realistic values (<1, 10, 30, 100, 300 μmol photons m^−2^ s^−1^, which equated to DLIs of <0.1, 0.3, 0.9, 3, 9 mol photons m^−2^ d^−1^), but more extreme combinations were also used to improve the model fit and assess the strength of the response.

#### Exp. 5. Artificial and realistic light spectra on A. tenuis settlement

A reduced set of experiments were conducted on *A. tenuis* in 2018 and 2019. The ‘realistic-broad’, ‘artificial blue, ‘artificial yellow, and ‘artificial red’ spectra (Experiment 1) were used in the 2018 experiment. And subset of treatment levels ‘realistic-broad’, ‘artificial red’ were conducted in 2019 to confirm results. Five simulated suspended sediment spectra (the same as Experiment 3) were used to investigate realistic suspended-sediment spectral effects.

#### Statistical analyses

All acute-exposure settlement data were fit to Bayesian binomial generalised linear mixed models, using a weakly informative prior (Cauchy; centre = 0, scale = 2.5) (Gelman et al. 2008) using the package *brms* (Bürkner 2017) in *R* (version 3.6.1). As hypotheses were planned, we did not remove any factors of interest during model selection even if not statistically significant, because all variables were of interest (Gelman & Hill 2006); however, comparisons of WAICs (Watanabe–Akaike Information Criterions) were used to assess if there were interactive effects. Standardised effect sizes and credible intervals of the posterior distributions were used to interpret the effect. Effect concentrations (EC10s), which are common effect-size threshold metric used in ecotoxicology, were calculated when possible (Warne et al. 2018).

### Settlement Series 2: Chronic exposure of water quality stressors to substrate quality

#### Exp. 6: Combined water quality stressors on substrate quality

To assess chronic exposure of water quality to the settlement cue, settlement discs were conditioned in the field under various light and sediment regimes. The discs were held in in PVC structures which orientated them either upwards facing or downwards facing. Each of the structures held 7 discs in different orientation and 3 structures were deployed at Florence Bay (relatively low turbidity; 19.121°S, 146.882°E) and Picnic Bay (relatively high turbidity; 19.186°S, 146.840°E) to condition for four months. Each structure was positioned randomly around a coral bombora to aid in retrieval in the turbid conditions. The upward facing surfaces are generally exposed to higher levels of light and accumulate deposited sediments, whereas downward facing surfaces are exposed to lower light intensities. One structure at each site included light loggers (Hobo Onset, MX2202) at each orientation to monitor relative light, calibrated to the MS8 multispectral sensor described above. Only the first few days of data after deployment were used because of algal fouling of the light logger sensor.

Approximately 1 week before the anticipated coral spawning, the structures were collected and transferred to flow-through aquaria in the SeaSim. One structure from Florence Bay was lost in turbid water during retrieval. The majority of macroalgae and sediment were removed from the discs as their inclusion would have made a census impossible. Settlement of larvae on these discs occurred in chambers under identical conditions using the methods described above in Settlement series 1 to remove any light or thigmotactic effects, and therefore the settlement response only represented changes in the substrate ‘quality’. Thirty competent *A. millepora* larvae were added to each experimental chamber under a 12:12 light/dark cycle for 24 hrs (30 μmol photons m^−2^ s^−1^; 1.3 DLI; broad-spectral profile) using the chamber set-up described above. Settled larvae were carefully counted using a Leica stereoscope by examining under all the remaining foliage. Discs were imaged with a Canon 90D camera using an EF-S 18–55 lens.

Settlement success on the discs were analysed with binomial GLMMs with ‘site’ and ‘orientation’ as fixed effects and ‘disc’ as a random effect with the package *lme4* (Bates et al. 2015). Pairwise comparisons were estimated with the package *emmeans* (Lenth et al. 2018). Community composition on the discs were analysed using photoQuad (v1.4) (Trygonis & Sini 2012), by grouping identifiable taxonomic and abiotic groups (macroalgae, green filamentous algae, red filamentous algae, CCA, cyanobacteria, tube worms, bryozoa, detritus, empty space, epilithic algal matrix (EAM), coral recruit, and unidentified). Taxonomic data was visually assessed using an nMDS (Bray-Curtis measure of dissimilarity) with the package *vegan* (Oksanen et al. 2007) using square-root transformed data. Environmental vectors and ellipses were subsequently added. To test differences in community structure between sites and orientation, PERMANOVA based on 999 permutations was implemented using the ‘*adonis’* function in *vegan*. Each distinct community was further examined to explore potential associations of taxa with settlement. Multicollinearity was assessed using variance inflation factors (VIF), pairs plots, and biological relevance (Zuur et al. 2010). Indicator Species Analysis using the package *labdsv* (Roberts & Roberts 2016) was first used to identify the most important taxa separating each group. Taxa with an indicator value >0.2 were used as a preliminary selection in a GLMM model. Corrected Akaike Information Criterion (AICc) was used to select the most parsimonious model.

#### Exp. 7: Sediment smothering on substrate settlement cue

While it was not feasible to investigate all water quality stressors on substrate quality, the effect of temporary sediment deposition on the settlement cue was examined as a likely mechanism (cause-effect pathway). Aragonite plugs (2-cm diameter) were conditioned for ~3 months in an outdoor aquarium system under 50% neutral density shade cloth (27 ° C; maxima 200 μmol photons m^−2^ s^−1^) and developed a thick covering of crustose algae (*Titanoderma prototypum* and *Peyssonnelia* spp.). The experiment aimed to simulate a sediment deposition and resuspension event and the plugs were exposed at various levels (0.1, 0.3, 0.8, 2.5, 8, 25, 83, 250 mg cm^−2^) of carbonate and siliciclastic deposited sediments (see Ricardo et al. (2018) for sediment characterisation) in flow-through conditions under ~30 μmol photons m^−2^ s^−1^ (PAR) linear-ramping lighting following Ricardo et al. (2017). The sediments were then removed, and each plug embedded in 60 mL glass chambers with cleaned river sand to prevent settlement on vertical or under-surfaces. Ten competent larvae were added to each chamber, and settlement on the sediment-free plugs was assessed at 24 hours. The plugs were then provided a recovery period of 3 days and then more larvae (n=10) were added to each chamber and settlement assessed after a second 24 hours. Effective quantum yields (ΔF/F_m_’), which estimates the photosystem II photosynthetic performance of light-acclimated samples, were measured on the plugs after a 30-minute light-acclimation period at 30 μmol photons m^−2^ s^−1^ with a mini-PAM (Waltz) using a red measuring light (6 mm fibre-optic probe) positioned ~ 3-mm away from the plug using a rubber spacer.

Measurements were taken before and after the exposure, and after the 3-day recovery period, with 5 measurements per plug. The changes in settlement were analysed with a binomial GLMM with ‘plug’ as random effect and the level of ‘previously-deposited sediment’ and time (after exposure, recovery) as a fixed factor. The changes in yield at each time point was measured with a four-parameter Weibull nonlinear model using the package *drc* (Ritz et al. 2016). The substrate composition was classified using Trainable Weka Segmentation in ImageJ (Arganda-Carreras et al. 2017) on a Dell R820 256 GB High-performance Computer System, and changes in proportion were predicted using a Dirichlet regression (Douma & Weedon 2019) with ‘previously-deposited sediment’ and time (before exposure, after exposure, and recovery) as a fixed factor.

## Results

### Settlement Series 1: Acute light exposure–experimental settlement trials

#### Exp. 1: Artificial light spectra on A. millepora settlement

The spectral profiles of monochromatic light treatments were generally eight-fold narrower compared to the broad-spectrum “white” treatment (Fig. 1 a). During the light phase, settlement under monochromatic blue light was similar to the control, and while there was a ~16% decrease in settlement under red light and a ~27% decrease under green light, there was a no statistically detectable difference (the 95% credible intervals did not overlap zero) (Fig. 1 b-c). There was a clear difference yellow light from the broad-spectrum control (~35% decrease) (Fig. 1 b-c). These trends indicate that coral settlement may be suppressed by up to one-third with wavelengths between the 500 and 600 nm range. Although there was a small amount of additional settlement in the subsequent dark period from 12 to 24 hrs (~ 25%), settlement remained suppressed in the green and yellow treatments indicating potential latency effects. The variance explained by the model was 72%.

**Fig. 1.**
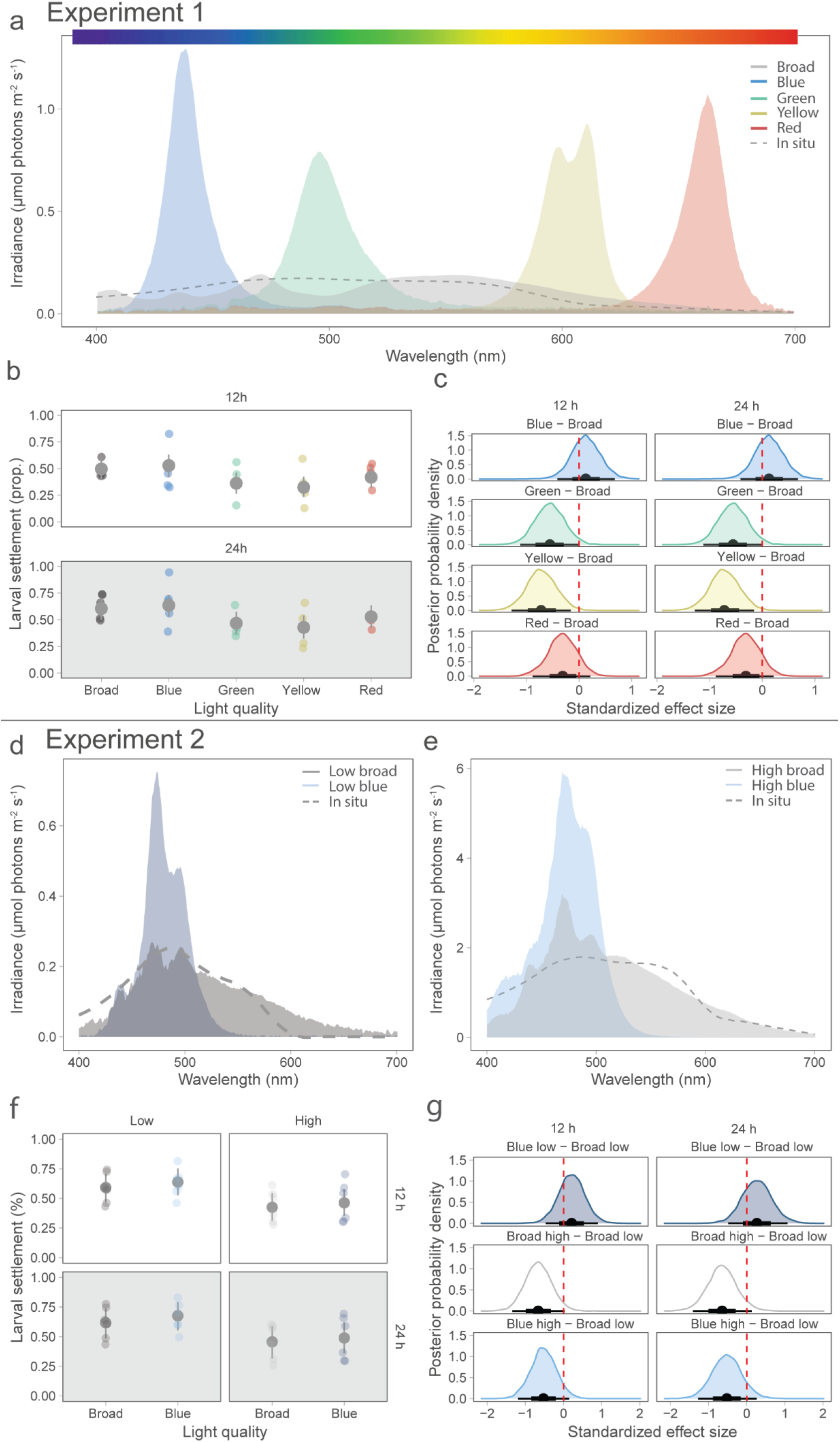
Larval settlement of *Acropora millepora* under artificial monochromatic and depth-dependant light spectral profiles in Exp. 1 & 2. (a) Light spectral profiles used in Exp. 1. Grey dashed line = shallow clear-water site at 5-m depth and 0.8 NTU. (b) Exp. 1 larval settlement after 12 h (light) and 24 h (12 light + 12 h dark) (light grey), modelled means and 95% credible intervals (dark grey). (c) Exp. 1 Standardised Bayesian posterior half-densities and credible intervals (thicker: 66%, thinner: 95%) of contrasts between each monochromatic light and the control (broad spectra). The vertical dashed red line indicates the x-axis location of zero effect. (d) Lower-light spectral profiles used in Exp. 2. Grey dashed line = deep clear-water site at 21 m depth and 1 NTU. (e) Higher-light spectral profiles used in Exp. 2. Grey dashed line = shallow clear-water site at 5-m depth and 0.8 NTU. (f) Exp. 2 settlement of larvae at two light intensities under broad-spectrum and artificial blue-light after 12 h (light) and 24 h (12 h light + 12 h dark). (g) Standardised Bayesian posterior half-densities and credible intervals (thicker: 66%, thinner: 95%) of the contrasts between each light spectrum and the control (broad low). The vertical dashed red line indicates the x-axis location of zero effect. Larval settlement competency was 90% for both experiments.

The light treatment wavelength in the choice experiment had generally wider spectral distributions than when each substrate was exposed independently because gel-filters were used instead of the multispectral lights (Fig. S1 a). There were no clear differences in settlement between the light treatments and broad-spectrum control, or over time (the 95% credible intervals did not overlap zero) (Fig S1 b-c). The variance explained by the model was 31%.

#### Exp. 2: Realistic depth-dependant light on A. millepora settlement

The low blue spectral profile had a narrower profile (FWHM = 39 nm) compared to the low broad profile (FWHM = 118 nm) (Fig. 1d). Likewise, the high blue spectral profile had a narrower profile (FWHM = 49 nm) compared to the low broad profile (FWHM = 81 nm) (Fig. 1e). There was a moderate (although not significantly detectable) 28% decrease in settlement under the higher light treatments (low: 0.62; 95% CI: 0.53–0.70); high: 0.44; 95% CI: 0.35–0.54), and no evidence of an effect of the blue dominant spectra on settlement (Fig. 1f-g). A further 12 hr dark phase did not increase settlement indicating no latency effects (Fig. 1f). The variance explained by the model was 78%.

#### Exp. 3: Realistic turbidity-dependant light spectra on A. millepora settlement

Low simulated-turbidity light treatments (~0.8–3 NTU) had similar spectral profiles whereas the highest simulated-turbidity profile (~9 NTU) was narrower and peaked around ~550 nm (Fig. 2a). The most parsimonious model only included ‘time’ as a predictor variable, indicating no evidence of an influence on settlement between the low turbidity and high turbidity-simulated spectra. Under the full model, there was no effect of the simulated turbidity spectra (slope = 0.1; 95% CI = −0.05– 0.25), and there was a slight overall increase of settlement between 12 hours (0.753, 95% CI: 0.721–0.783) and 24 hours (0.790, 95% CI: 0.761–0.819) (Fig. 2b,c). The variance explained by the full model was 25%.

**Fig. 2.**
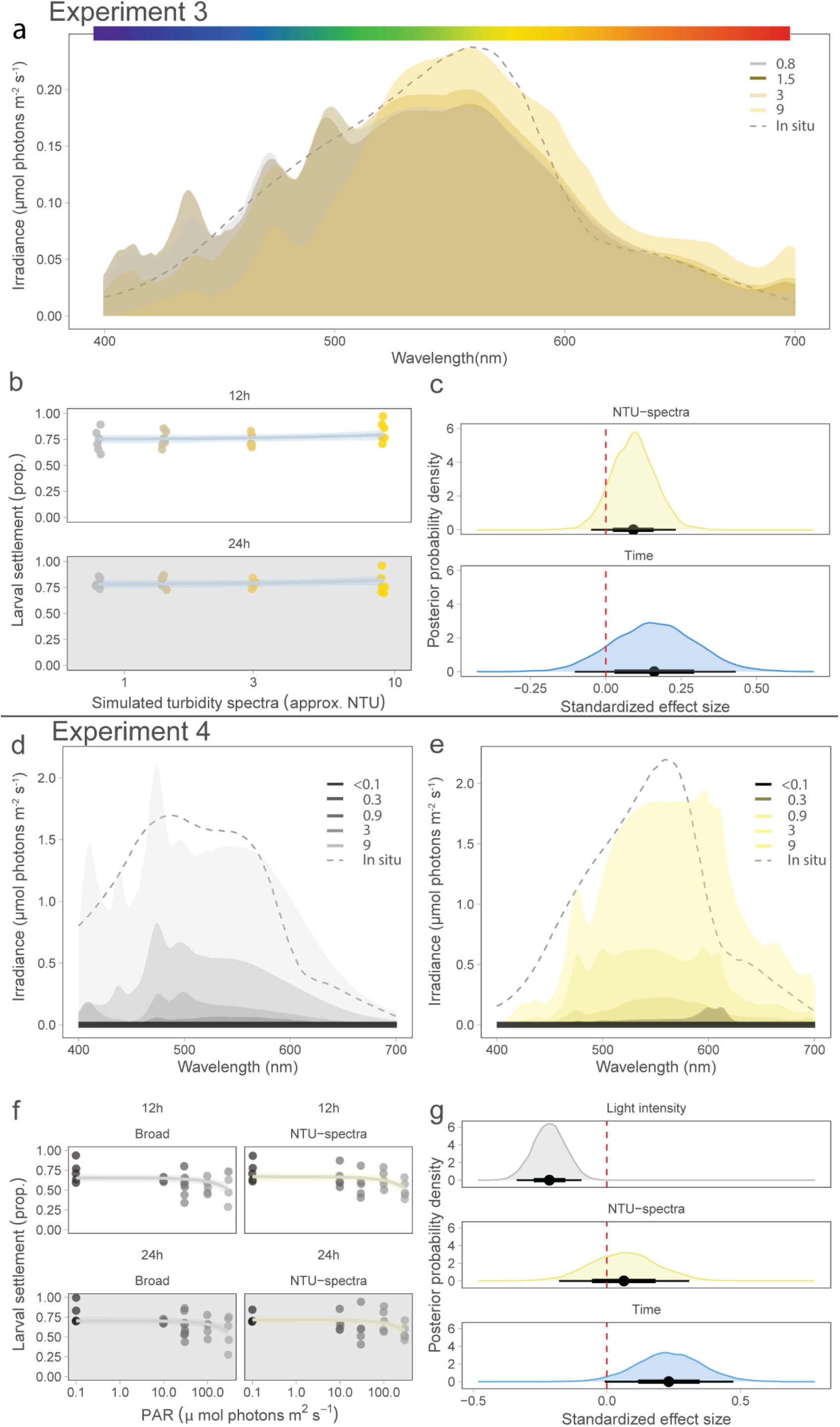
The relationship between light intensity and simulated turbidity spectra on larval settlement of *Acropora millepora*. (a) Exp. 3 spectral profiles used to simulate in situ spectral patterns at various approximate turbidity (NTU) levels (Jones et al. 2020). Grey dashed line = shallow clear-water site at 5-m depth and 0.8 NTU. (b) Exp. 3 larval settlement in the turbidity spectral treatments after 12 h (light) and 24 h (12 light + 12 h darkness). (c) Exp. 3 standardised Bayesian posterior half-densities and credible intervals (thicker: 66%, thinner: 95%) of contrasts. The vertical dashed red line indicates the x-axis location of zero effect. (d) Exp. 4 spectra used to simulate clear-water spectral patterns at five light intensities. Grey dashed line = shallow turbid-water site at 5-m depth and 5 NTU. (e) Exp. 4 spectra used to simulate high turbidity spectral patterns at five light intensities. (f) Exp. 4 larval settlement after 12 (light) and 24 hours (12 h light + 12 h dark) of the light exposure. (g) Exp. 4 standardised Bayesian posterior half-densities and credible intervals (thicker: 66%, thinner: 95%) of the contrasts. The vertical dashed red line indicates the x-axis location of zero effect. Larval settlement competency was 90% in both experiments.

#### Exp. 4: Light intensity in clear and turbid spectra on A. millepora settlement

The clear-water spectral profiles generally had a peak at in the blue wavelength (~480 nm) whereas the turbidity spectral profiles peaked in the green wavelengths (~550 nm) (Fig. 2d,e). There was no difference in settlement between the clear and turbid spectra treatments but a statistically detectable (the 95% credible intervals did not overlap zero) modest 22% decline in settlement at higher light intensities (12 hr: EC10 = 113 μmol photons m^−2^ s^−1^; 95% CI: 11– 214) (Fig. 2f,g). PAR measurements at the clearer-water site (Florence Bay) exceeded this threshold 21.2% of the time. There was also a slight overall improvement in settlement of 8.4% following the dark-period, but there remained evidence of latency effects at higher light intensities (Fig. 2f-g). The variance explained by the model was 66%.

#### Exp. 5. Artificial and realistic light spectra on A. tenuis settlement

Settlement of *A. tenuis* in the broad spectra was low in both years (2018: ~20%; 2019: ~37%) compared with the settlement competency assays (>80%) indicating sub-optimal conditioning (CCA and biofilm) of the substrates for this species (Fig. 3a,c,e). In both years, the artificial red light increased settlement (2018: 2.2-fold greater, R^2^ = 0.87; 2019: 1.4-fold greater, R^2^ = 0.88) (Fig. 3a-d). There was no effect of the realistic dredge spectra (yellow-green dominant) on coral settlement (Fig. 3e,f), with 84% of the variance explained by the model.

**Fig. 3.**
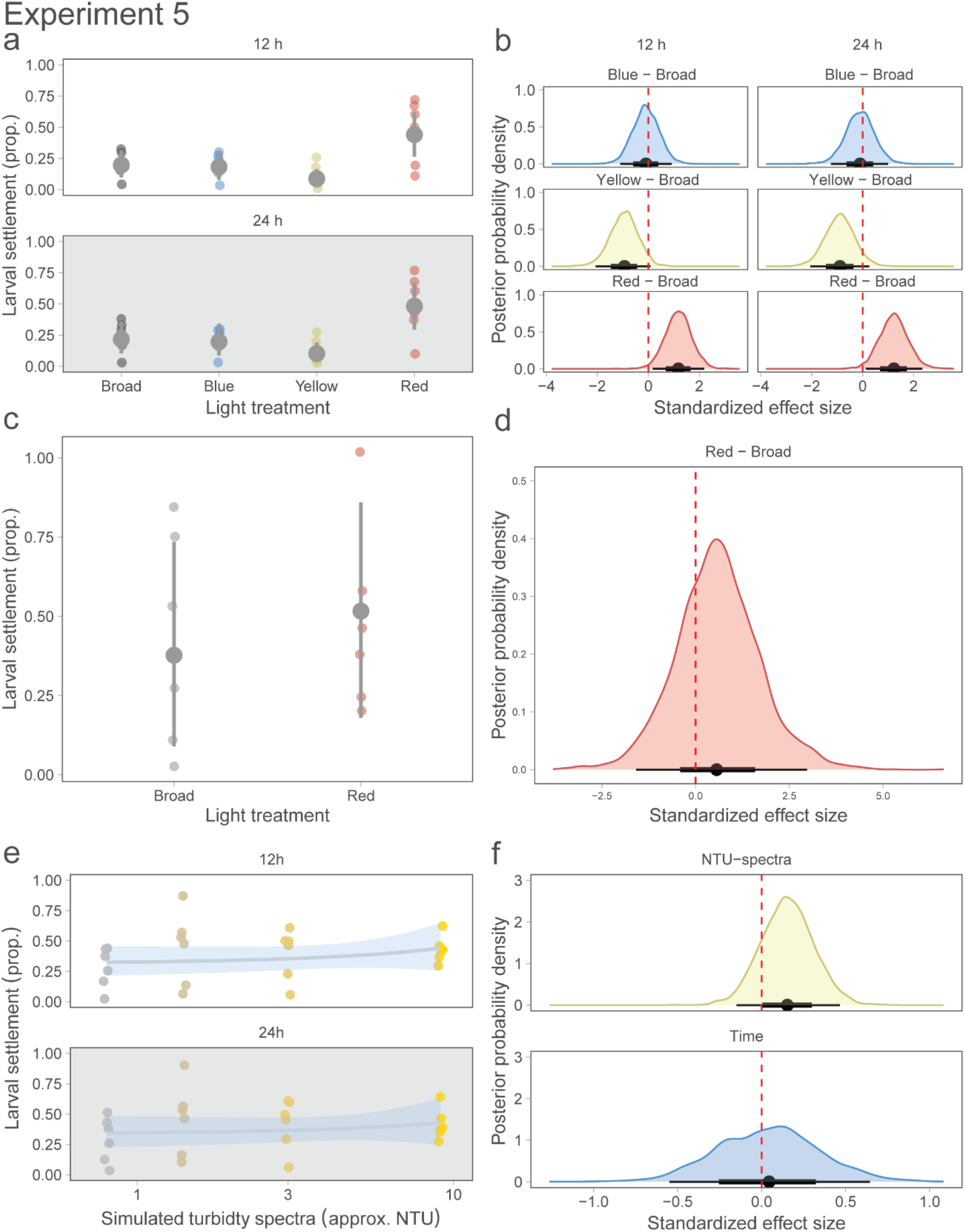
Effects of artificial (monochromatic) and simulated turbidity spectral profiles on larval settlement of *Acropora tenuis* in Exp. 5. (a) Settlement after 12 and 24 hrs. Spectral profiles matched Exp. 1 (broad, artificial blue, artificial yellow, and artificial red). (b) Standardised Bayesian posterior half-densities and credible intervals (thicker: 66%, thinner: 95%) of contrasts between each monochromatic light and the control (broad spectra). The vertical dashed red line indicates the x-axis location of zero effect. (c) 2019 repeat experiment at only the broad and artificial red treatments. (d) Standardised Bayesian posterior half-densities and credible intervals (thicker: 66%, thinner: 95%) of contrasts between each red light and the control (broad spectra). (e) Settlement in simulated-turbidity spectra reported in NTU after 12 h (light) and 24 h (12 light + 12 h darkness). (f) Standardised Bayesian posterior half-densities and credible intervals (thicker: 66%, thinner: 95%) of contrasts. Larval settlement competency for all assays were >80%.

### Settlement Series 2: Chronic exposure of water quality stressors to substrate quality

#### Exp. 6: Combined water quality stressors on substrate quality

##### Water quality

The light environments of downwards-facing surfaces were 8–9 fold lower than that of upwards-facing surfaces on the field-deployed discs (Table 3). Light intensity was 30% lower on the upward-facing surfaces at the clearer-water site (Florence Bay) compared with the higher-turbidity site (Picnic Bay), and 20% lower on the downward-facing surfaces.

**Table 3.** Water quality at the clearer (Florence Bay) and turbid (Picnic Bay) water sites and each substrate orientation. Light was measured during the first 3 d of deployment. NTU was measured for 1–2 mo. depending on the site.

| Site | Orientation | DLI | NTU |
| --- | --- | --- | --- |
| Florence Bay | Upwards | 18.5 (n=2) | 0.6 |
| Florence Bay | Downwards | 2.07 (n=2) |  |
| Picnic Bay | Upwards | 12.9 (n = 2) | 1.62 |
| Picnic Bay | Downwards | 1.65 (n = 1) |  |

Turbidity, which can be approximately converted to suspended sediment concentrations using a conversion factor of 1.1 (Jones et al. 2020), was greater at Picnic Bay (50^th^ percentile = 1.62 NTU) compared to Florence Bay (50^th^ percentile = 0.60 NTU).

##### Settlement

There was a clear difference in settlement on field conditioned discs at each orientation, with the effects more pronounced at the clearer-water site (Florence Bay), compared with the more turbid-water site (Picnic Bay) (Fig. 4a,b). At Florence Bay there was a ~74% decrease (>99.99% probability of a negative effect) in settlement on the upwards-facing discs relative to the downward-facing discs. A similar although more modest trend occurred at Picnic Bay with a ~51% decrease (99.91% probability of a negative effect) in settlement on discs conditioned upwards. However, settlement on downwards-facing surfaces was lower at on discs conditioned at the more turbid Picnic Bay site by 36%. The variance explained by the model was 87%.

**Fig. 4.**
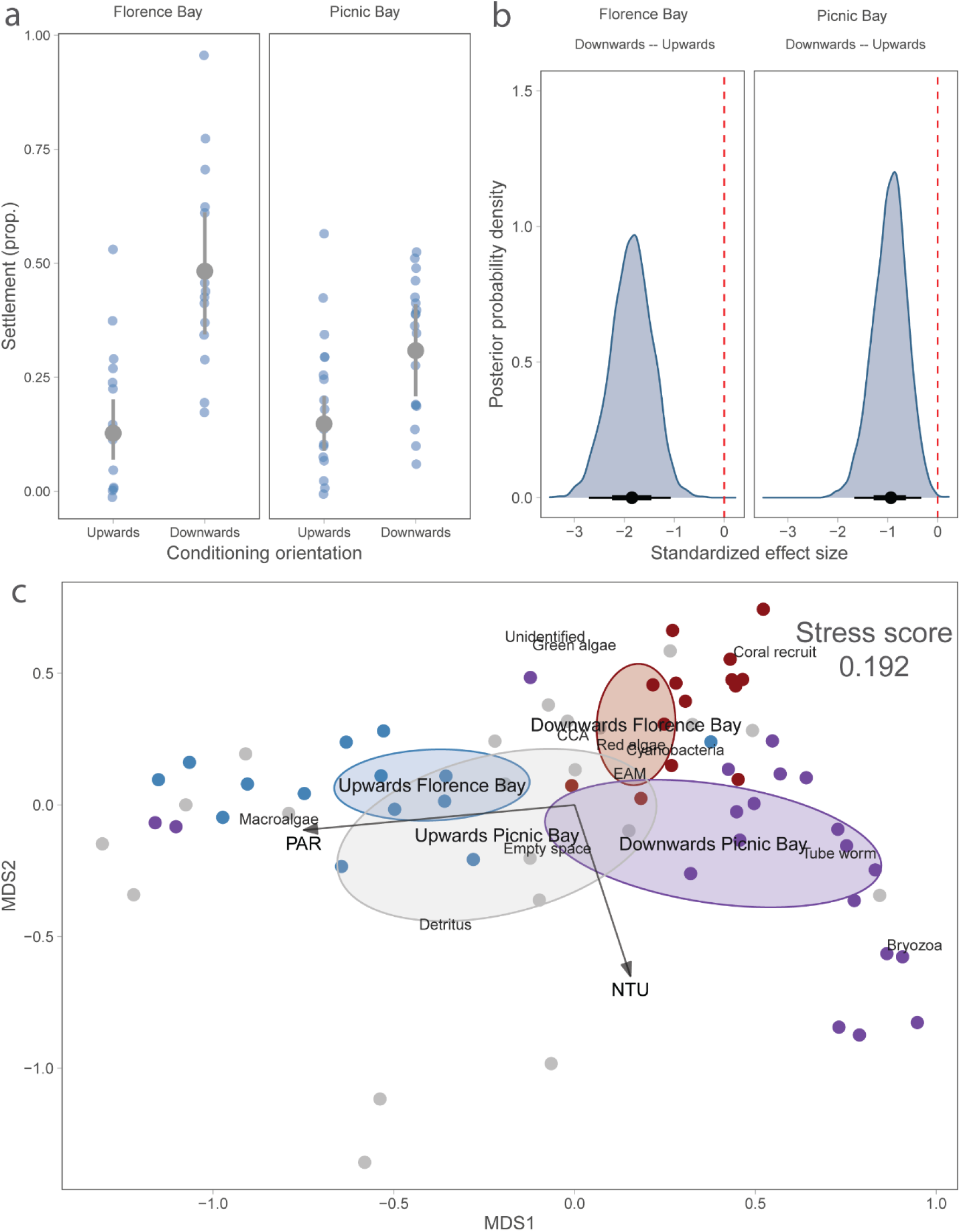
Settlement and community composition of the settlement discs in Exp. 6. Settlement on the discs conditioned at (a) Florence Bay (clearer-water site) and (b) Picnic Bay (turbid-water site). All settlement occurred with discs facing upwards and under constant-light conditions. Larval competency was 80%. (c) An nMDS plot showing community composition at each of the site and orientation combinations. Ellipses represent 1 standard deviation around the centroids. Vectors represent the environmental factor fit. CCA = crustose coralline algae; EAM = Epilithic algal matrix, PAR = photosynthetically active radiation; NTU = Nephelometric Turbidity Units.

The community composition was different on discs at each site and orientation (Fig. 4c; PERMANOVA, p = 0.002), with clear separation between the upwards and downwards facing with discs. For Florence Bay downwards-facing discs, there was a negative correlation between crustose coralline algae and other red algae (r = −0.74, VIF = 2.78), consequently ‘red algae’ was subsequently removed from the analysis. The exploratory analysis revealed

CCA was associated with Florence Bay downwards-facing discs, tube worms with downwards-facing surfaces at Picnic Bay, and empty space (uncolonized by macro-organisms) on Picnic Bay upwards-facing surfaces (Fig. S2).

#### Exp. 7: Sediment smothering on the substrate settlement cue

##### Settlement

After the 3-day exposure to carbonate sediment, there was a clear decrease in settlement on the substrates exposed to the greatest levels of deposited sediment (Fig. 5a, EC10 = 38 mg cm^−2^; 95% CI: 8.5–68). There was a slight improvement in larval settlement on the same substrate after the 3-day recovery period (Fig. 5c, EC10 = 63 mg cm^−2^; 95% CI: 0–134). A similar yet less extreme response was observed following exposure to siliciclastic sediment (Fig. 5b, EC10 = 64 mg cm^−2^; 95% CI: 16–112), with less influence on settlement after the recovery period (Fig. 5d, EC10 = 90 mg cm^−2^; 95% CI: 0–217).

**Fig. 5.**
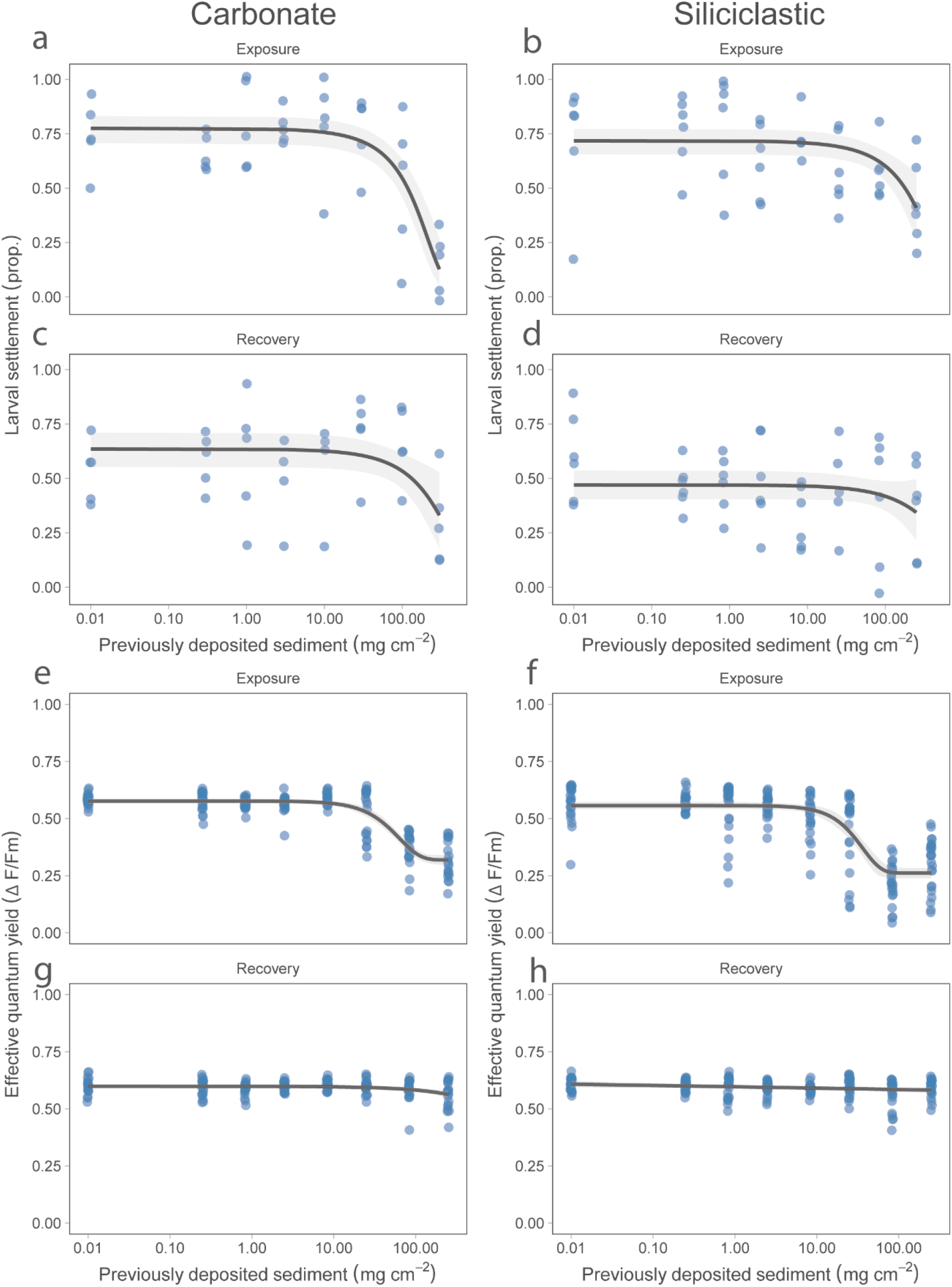
Settlement of *Acropora millepora* larvae on CCA previously covered for 3-days in two types of deposited sediment in Exp. 7. (a) Settlement following removal of carbonate sediment. (b) Settlement following removal of siliciclastic sediment. (c, d) Settlement following a 3-day recovery period for each sediment type. (e) Effective quantum yields (ΔF/F_m_’) of CCA following removal of carbonate sediment (f) ΔF/F_m_’ following removal of siliciclastic sediment. (g, h) ΔF/F_m_’ following a 3-day recovery period for each sediment type. Larval settlement competency was 80%.

##### Sub-lethal indicators

Effective quantum yields (ΔF/F_m_’) of the CCA on the settlement substrate decreased within the same effect region (Fig. 5e,f; carbonate: EC10 = 28; siliciclastic: EC10 = 16 mg cm^−2^) as larval settlement success, and also recovered after a 3-day period (Fig. 5g,h; EC10s >250 mg cm^−2^). Before exposure, CCA of *Titanoderma prototypum* covered ~61% of the plugs, and *Peyssonnelia* spp. covered ~21% of the plugs. Bleaching (mostly driven by impacts on *T. prototypum*) increased proportionally with sediment smothering (evident at > 30 mg cm^−2^) and did not regain pigment after the 3-day recovery period (Fig. S3a-c). This indicates that changes in ΔF/F_m_’ were a better predictor of settlement cue effectiveness than observed bleaching.

## General Discussion

Disentangling the various drivers affecting coral recruitment that occur sequentially and simultaneously remains a challenge of marine ecology (Graham 2003, Houlahan et al. 2017). While little can be done to control all co-occurring factors in observational studies (Graham 2003), manipulative studies offer unique opportunities to identify and quantify key drivers that influence coral recruitment patterns and other biological processes (Chapman 2016, Nosek & Errington 2020). For example, a number of correlating variables have been described to predict coral spawning events, yet careful manipulation of only two (temperature and light intensity) are required to reliably control the timing of spawning *ex situ* (Craggs et al. 2017). Here we sought to test the relative importance of direct ‘acute’ light compared to chronic water-quality factors impacting the substrate by applying highly controlled but realistic experimental approaches. We first characterised the light environment at scales relevant to settling larvae using spectroradiometers, and then used these data to inform a selection of experimental treatment levels using a multispectral feedback LED lighting system. In the case of *Acropora*, only unrealistic artificially narrow or high light-intensity exposures led to settlement shifts in comparison to the lower broad-spectrum control light. For example, monochromatic light of mid-range (green-yellow) wavelengths influenced settlement negatively in some experiments; however, these effects were not observed under a broader realistic spectra that occur in shallow waters, at depth and under high turbidity. To improve the likelihood of detecting effects of light spectrum and intensity on settlement, we opted for a sub-optimal conditioning to the discs in comparison to Ricardo et al. (2017) where a more mature substrate community dominated by CCA may have overwhelmed light-intensity effects. Nevertheless, the present study similarly indicated high light-intensity has only a modest negative effect on settlement at levels typically found during midday on exposed surfaces. It is not entirely surprising that larvae searching the benthos are relatively insensitive to light intensity, as strong light gradients are only available for a few hours each day. Interestingly, *A. tenuis* did show positive effects to artificial red light, especially in one spawning year. In these experiments the larvae had high competency (ability to settle) but low settlement success in the control treatments indicating sub-optimal conditioning, and this supports the possibility that weaker settlement inducers/inhibitors (such as direct light) may become more important in absence of strong settlement inducers such as mature CCA colonies.

Several lines of evidence suggest that water-quality conditions acting on the substrate prior to settlement may be critical for determining coral settlement outcomes (Baird et al. 2003, Petersen et al. 2005, Kuffner et al. 2006, Szmant & Miller 2006, Suzuki & Hayashibara 2011, Ricardo et al. 2017). Chronically altered light regimes and smothering by sediment lead to altered community assemblages on the substrate that evidently promote or prevent settlement (Erwin et al. 2008, Arnold & Steneck 2011, Quimpo et al. 2020). In this study, prior exposure of the settlement substrates to a variety of water quality conditions *in situ* elicited clear differences in the settlement response, with larvae preferring the substrates that had been previously conditioned under low light and not affected by sediment. While there remains some limitations in characterising the community on settlement substrates visually and inferring casual effects in absence of measuring possible confounders (Walker 2014), CCA colonised downward-facing surfaces at the clear-water site was associated with high settlement, whereas tubeworms were associated with decreasing settlement on the downwards-facing surfaces conditioned at the turbid-water site. Empty space (possibly grazed clear by gastropods observed at the time of retrieval) was associated with high settlement on the upwards-facing surfaces. Known biological settlement inducers such as CCA are often associated with specific light environments (Petersen et al. 2005, Bessell-Browne et al. 2017, Briggs & Carpenter 2019) but are sensitive to sediment smothering (Harrington et al. 2005, Ricardo et al. 2017), and searching larvae may favour these communities that could be advantageous for post-settlement growth. However, optimal conditions and responses of biological inducers for coral settlement (CCA and microbial biofilms) to stress remain remarkably understudied (Webster et al. 2011, Webster et al. 2013), and derivation of water quality and light thresholds for important algal and microbial inducers would improve predictions for coral settlement.

In addition to the potential for chronic effects of light exposure on the benthic surfaces to influence its structure and suitability for coral settlement, highly turbid conditions are also associated with sediment deposition (Whinney et al. 2017), which can accumulate on settlement surfaces directly affecting access by larvae (Hodgson 1990, Ricardo et al. 2017), or indirectly affecting settlement by harming biological settlement inducers such as CCA (Kendrick 1991, Harrington et al. 2005). In the current study, there were strong impacts of sediment accumulation and declining quality of settlement cues, resulting in a subsequent decrease in coral settlement. Even brief sediment exposures to the substrate at levels that are common for inshore reefs (see Tebbett et al. (2018), and Jones et al. (2020)) led to a decrease in settlement success, as well as physiological responses of the substrate such as pigment loss (bleaching) and decreased ΔF/F_m_’. Sediment smothering can reduce gas exchange, prevent waste removal, eliminate light, and change the surface chemistry of the smothered organism (Weber et al. 2012, Jones et al. 2016). The settlement response we observed was more pronounced with carbonate sediment, and the differences between the two sediment types is possibly caused by seemingly rapid consolidation of these sediments over the 3-day period, with the carbonate sediment bound more firmly to the substrate at the time of removal. Following the subsequent 3-day recovery period, ΔF/F_m_’ and the dose-dependent settlement increased, whereas bleaching did not change, indicating photo-physiological responses maybe more effective indicators of settlement success than bleaching (loss of pigments). Although some recovery was observed following sediment removal, turbid reefs are subject to ongoing sediment deposition and removal processes that likely result in eventual loss or reduction of settlement inducers on upwards-facing surfaces over longer durations (Kendrick 1991).

It is often difficult to interpret studies assessing impacts of light or other water-quality factors during coral settlement where substrates were not randomised immediately before settlement, and where surfaces vary in sizes, roughness and aspects, introducing the possibility of edge and orientational effects (Raimondi & Morse 2000). This is common in field settlement assays where typically ~10 × 10 × 1 cm ceramic tiles are conditioned *in situ* for a few weeks prior to the mass spawning events (Table S1), and there have been calls for greater standardisation in settlement studies (Abelson & Gaines 2005, Davidson et al. 2019). Further, larvae exhibit strong thigmotactic behaviour with an attraction to grooves, edges and aspects in the absence of light (Nozawa et al. 2011, Whalan et al. 2015, Ricardo et al. 2017).

Edmunds et al. (2014) and Nozawa et al. (2011) suggests that a reduction in settlement on horizontal surfaces of field-deployed settlement tiles may be due to a lack of micro-refuges. However for inshore reefs, deposited sediment and algal mats may infill these refuges driving settlement to vertical and downwards-facing surfaces (Ricardo et al. 2017, Tebbett & Bellwood 2019). Importantly, downwelling irradiance is confounded by orientation and edge effects because of local shading, leading some authors to speculate that direct light intensity is the driver of coral recruits settling near the edges of undersides (Morse et al. 1988, Maida et al. 1994). The current study removed confounding thigmotactic and orientation effects that are common on three-dimensional substrates. Further crossed designs with randomisation of the substrate prior to settlement are needed to conclusively separate light-intensity effects from possible thigmotactic and substrate effects.

For the species tested here, and possibly many species in the genus *Acropora*, the influence of light and physical challenges (such as sediment deposition and algal growth) on settlement of the larvae is predominantly manifested indirectly by influencing the biology and quality of settlement surfaces, rather than light as a direct cue influencing the settlement response.

However, in some species, there is greater ‘weight of evidence’ supporting the idea of light sensitivity in settlement in other genera such as *Oxypora* (Babcock and Mundy, 1996; Mundy and Babcock, 1998). Perhaps a more plausible explanation of settlement patterns under various narrow-band spectra is a mistaken recognition of surface colour (i.e., spectral diffuse reflection) from specular reflection. Several studies have observed increased settlement on orange-red surfaces and have suggested there is an attraction because CCA species typically have a reflectance peak in the 600 nm region of the visible spectrum (Mason et al. 2011, Foster & Gilmour 2016). Alternatively, the attraction may be related to the loss of blue light reflected from these surfaces, as reduced blue light is associated with temporary reduced swimming speeds in larvae (Sakai et al. 2020). In our study, reduced settlement was observed on CCA that had lost its red pigments (bleached), but after a recovery period improved settlement correlated well with recovered photosystem activity, suggesting that physiological processes (an indicator of healthy CCA) were a stronger driver of settlement than spectral reflectance. Other studies have reported phototaxis of coral larvae in the water column (Lewis 1974, Szmant & Meadows 2006, Mulla et al. 2020, Sakai et al. 2020), although these behaviours appear less consistent than observed in other marine invertebrates (Raimondi & Morse 2000, Satheesh & Wesley 2010, Ricardo et al. 2014, Lillis et al. 2018, Abdul Wahab et al. 2019).

The findings presented here suggest targeted management of water quality (e.g., sediment plumes from dredging) in the weeks to months leading up to coral spawning events may be a more meaningful area of focus for management rather than solely focussing on managing water quality during the days of predicted settlement. Deposited sediments are known to strongly repel coral settlers on upwards-facing surfaces (Ricardo et al. 2017), and while the temporal dynamics (i.e. deposition vs resuspension) remains unresolved, evidence is building that acute deposition events strongly tied to anthropogenic activities such as dredging can be managed with a spatial separation area of a few kilometres (Jones et al. 2020). However little evidence suggest that a short-term cessation of close-proximity dredging would improve settlement outcomes, as the sediment would need time for resuspension, latent effects on the substrate would remain for several days, and the chronic change in the water quality environment would cause shifts in the substrate community. Likewise, improvements in water quality and other factors that improve the substrate for settlement such as fish herbivory (to control algal growth) would likely improve settlement success (Doropoulos et al. 2016, Tebbett & Bellwood 2019).

Here we show for two common *Acropora* species that realistic light quality and quantity at the time of settlement plays a lesser role in determining settlement success in comparison to chronic water quality conditions impacting the settlement substrate over weeks to months prior to settlement – contrary to expectations. Teasing apart confounding factors inherent in settlement experiments will require innovative and creative experimental approaches, and with advancements in technology and better characterisation for the settlement environment, there are opportunities to further test, refine and reject old ideas. As researchers race to find ways to increase local reef resilience and scale-up restoration via seeding of coral larvae (Randall et al. 2020), it is crucially important to understand which factors primarily drive recruitment, and which act in a more subtle or infrequent manner.

## Supporting information

Highlights

Graphical Abstract

## Data accessibility

All raw data used in the figures can be accessed here https://github.com/gerard-ricardo/light-settlement.

## Acknowledgements

This project was supported through government appropriation funding (AIMS) and from the Australian Government’s National Environmental Science Program (NESP). We acknowledge In situ Marine Optics for their technical expertise with the light sensors. We acknowledge Eduardo Arias, Matt Salmon, Paul Boyd, Andrea Severati for design of the sediment dosing and lighting systems, and the AIMS SeaSim team for assistance for providing conditioning tanks, room set-up and animal husbandry requirements. We would also like to thank Lee Bastin, Florita Flores, Marie Thomas, Chris Brunner, Annika Lamb, Lonidas Koukoumaftsis and Christian Odea for assistance with coral spawning and image analyses.

## Notes

### Competing Interest Statement

The authors have declared no competing interest.

### Summary of Updates

Added Highlights and Graphical Abstract. Minor edits.

