## Supplementary Material for "Impacts of water quality on *Acropora* coral settlement: The relative importance of substrate quality and light"

### Text S1.

The genus *Acropora* is a major framework builder and coloniser on the Great Barrier Reef (GBR) and the Indo-Pacific (Sheppard et al. 2008, Hughes et al. 2019), and two species (*Acropora millepora* (Ehrenberg, 1834) and *Acropora tenuis* (Dana, 1846)) were selected to represent this genera. Gravid adult colonies were collected from <6 m depth from inshore reefs of the central GBR (*A. millepora*: Falcon Island: 18.766°S, 146.533°E; *A. tenuis*: Magnetic Island: 19.104°S, 146.862°E) and transported to the National Sea Simulator (SeaSim) at the Australian Institute of Marine Science (AIMS), Townsville in the later months of 2018 and 2019. On the nights of spawning, egg-sperm bundles from six to eight colonies were collected and cross fertilised. Embryos were then washed free of sperm and transferred into 500 L flow-through fiberglass tanks to undergo larval development following the methodologies of Heyward and Negri (1999) and Ricardo et al. (2018).

Only larvae that had a settlement competency of least 70% and did not 'self-settle' were used. Low levels of settlement competency in assays could indicate an unhealthy larval culture and therefore larvae could be overly sensitive to stress (EC 2005, Guest et al. 2010). Alternatively 'self-settlement' (settlement without a settlement inducer) could indicate overly competent larvae that may be insensitive to the treatment. Settlement competency assays involved testing for settlement success of the larval culture before commencement of the experiments in 6-well plates by providing ten larvae (2–3 wells) to ~4-$mm^2$ live chips of *P. onkodes* in FSW, and one well (n = 10 larvae) without an settlement inducer designated as a negative control (Ricardo et al. 2016). All assays were conducted over 24 hrs in room light (~10 µmol photons $m^{-2}$ $s^{-1}$, 10:14 light/dark cycle).

### Text S2.

In-situ measurements of this study were conducted on the reefs or waters around Magnetic Island, Cleveland Bay in the inner central Great Barrier Reef (GBR). The Bay is naturally turbid with natural wind-wave conditions often sufficient to resuspend bottom sediments at 5 m depth (Orpin & Ridd 2012, Macdonald et al. 2013, Whinney et al. 2017). Turbidity generally decreases towards the head of the Bay, producing a north-east turbidity gradient (Jones et al. 2020). Maintenance dredging operations that increase turbidity are also common along the eastern side of Magnetic Island (Pringle 1989, Jones et al. 2020). Sediments visually appear brown and are primarily composed of a mix of terrestrial (siliciclastics) and biogenic (carbonates) minerals (Duckworth et al. 2017).

Before experiments commenced, a light meter connected to a diving pulse amplitude modulating fluorometer (PAM) (Walz) calibrated to a LI-COR (LI-250A) quantum light meter was used to measure instantaneous light intensities (PAR) in ~10 × 10 cm crevices along the substrate at Middle Reef (19.196°S, 146.814°E) in the inshore ‘turbid reef zone’ (Browne et al. 2010), at 3–5 m depth during midday in November 2018 (Table 2). Coral larvae have a tendency to settle in areas not directly in light such as grooves, under algae and on vertical and undersides (Petersen et al. 2005, Nozawa et al. 2011, Davidson et al. 2019), and these measurements were used to determine the environmentally realistic range of light intensities that coral recruits are likely to settle and grow *in situ* (but see Discussion below). Mean instantaneous PAR levels were 15.6 µmol photons $m^{-2}$ $s^{-1}$, magnitudes lower than that typically occur on shallow-exposed-horizontal surfaces (Jones et al. 2020). We conservatively selected 30 µmol photons $m^{-2}$ $s^{-1}$ for the control light intensity in experiments.

Two continuous-measurement (every 15 min) MS8 multispectral radiometers (In Situ Marine Optics, Perth, Australia) with irradiance measured at eight predetermined wavelengths (~30 nm increments) were deployed in ~5-m depth at Florence Bay (lower turbidity; 19.121°S,

146.882°E) and Picnic Bay (high turbidity; 19.186°S, 146.840°E) reefs to measure temporal light-intensity percentiles and approximate spectral profiles under turbidity. These data were used to inform simulated light-intensity treatments for use in laboratory experiments and spectral profiles at various turbidity.

In September 2016, light spectral profiles were measured in and around Cleveland Bay, Townsville using a USSIMO hyperspectral radiometer (In Situ Marine Optics, Perth, Australia) with irradiance (W $m^{-2}$ $nm^{-1}$) measured at 5-nm bands between 400 and 700 nm. At the furthest offshore clear-water site (19.085°S, 146.947°E) a light spectral profile was measured each at 5 m and 21 m depth. Another profile was measured at a highly turbid-water site at 5 m depth (19.169°S, 146.905°E). These data were used to inform spectral profiles across the simulated depth and turbidity light gradients used in laboratory experiments. The light spectral profile at 5-m depth (~0.8 NTU) was applied as the experimental control in all experiments. Additionally, individual light channels were used to replicate monochromatic wavelengths commonly applied in spectral-light response studies.

### Text S3. The influence of monochromatic light spectra on settlement choice

To assess whether subtle effects of spectral quality affected settlement of *A. millepora*, larvae were presented with a choice of substrates in close proximity (~4 cm) to each other, but exposed to various spectra: broad spectrum ($\lambda$ = 570 nm; 162nm), blue ($\lambda$ = 452 nm; 22 nm), green ($\lambda$ = 547 nm; 64 nm), yellow ($\lambda$ = 588 nm; 115 nm), red ($\lambda$ = 638 nm; 73 nm), and five Hydra FiftyTwo HD™ (AquaIllumination Inc.) aquarium lights were positioned above coloured gel filters (Rosco). The light intensity was controlled at 29 ± 3 µmol photons $m^{-2}$ $s^{-1}$ (mean ± SD) PAR under a 12:12 light/dark profile. Each disc was slotted into a 24 cm diameter circular unconditioned PVC canister that sat within a settlement chamber (a glazed ceramic dish). Teflon tape was wrapped around the circumference of the PVC canister to prevent settlement on vertical surfaces and cleaned river-sand was used to prevent larvae entering under the canister. One hundred larvae were pipetted into the centre of the dish, and settlement was assessed at 12 and 24 h as described above.

### Fig. S1. The influence of monochromatic light spectra on settlement choice

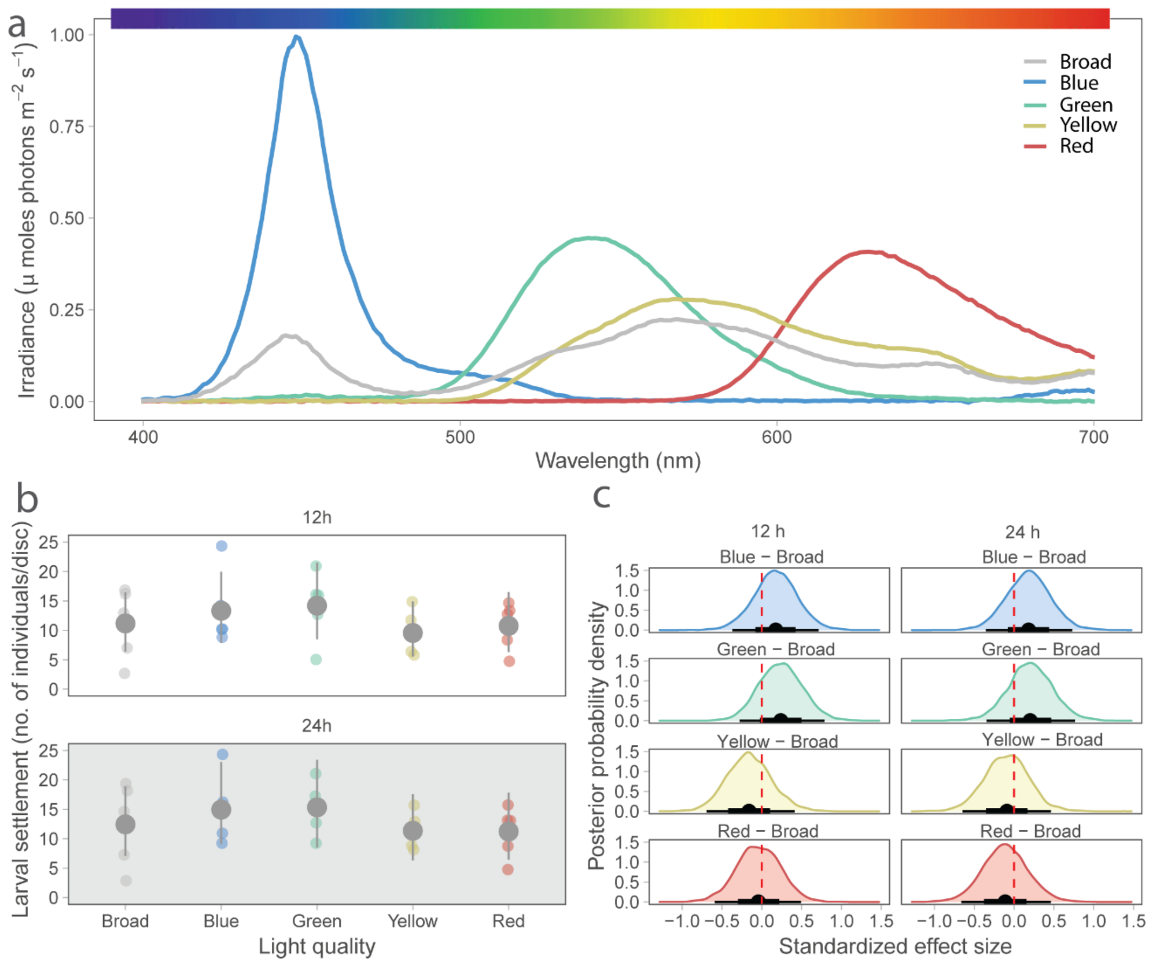


Figure S1. Larval settlement of *Acropora millepora* under artificial and broad-spectrum control lighting using a ‘choice’ design in Exp. 1. a) Spectral profiles of the five light treatments binned to the nearest nanometre. b) Mean larval settlers per coloured disc and 95% credible intervals after 12 hours and 24 hours exposure to light regimes. (c) Standardised Bayesian posterior half-densities and credible intervals (thicker: 66%, thinner: 95%) of contrasts between each monochromatic light and the control (broad spectrum). The vertical dashed red line indicates the x-axis location of zero effect. Larval settlement competency was 80%.

Fig. S2 Community composition for field conditioned discs

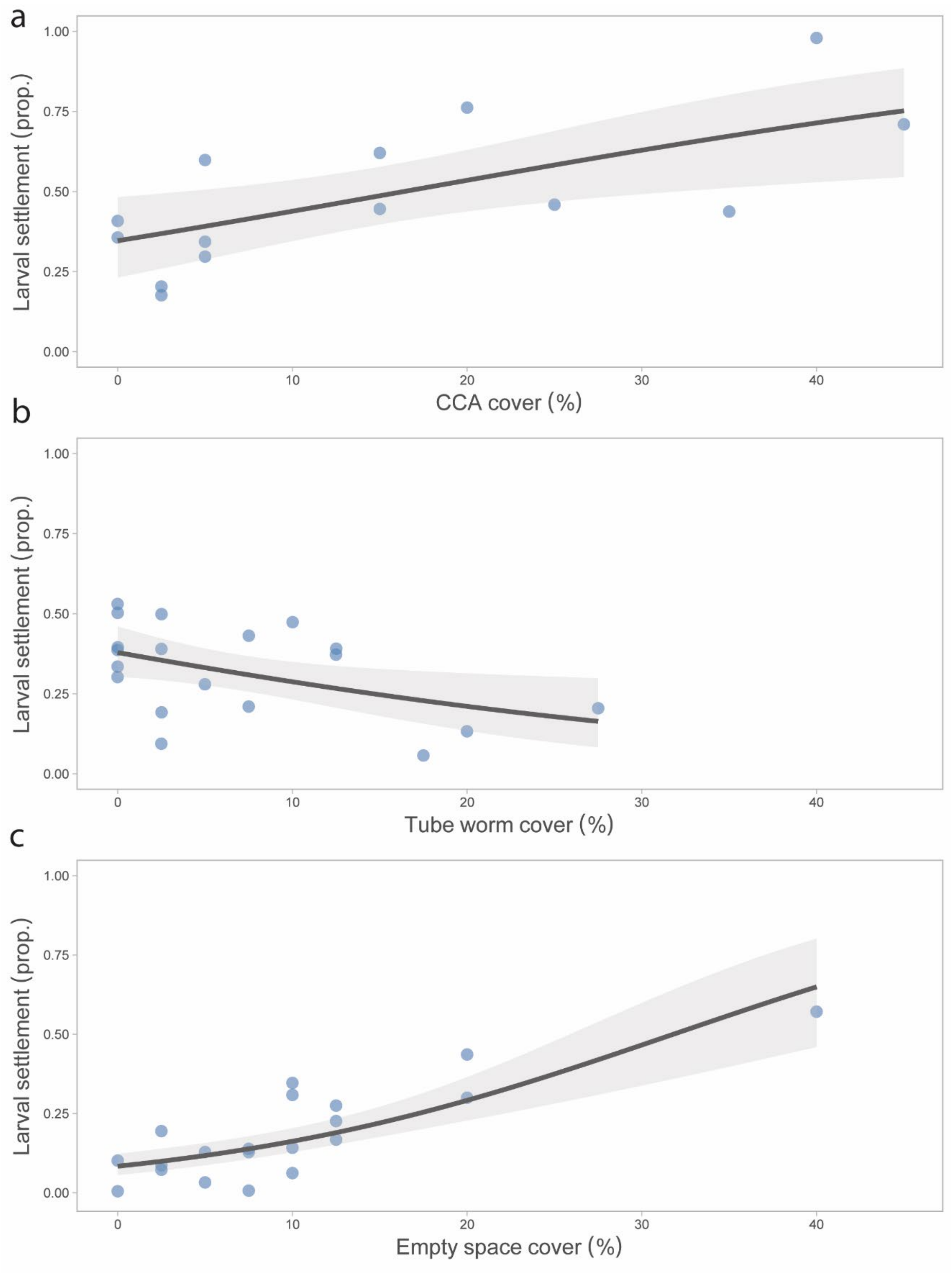

Fig. S2. Partial regression plots between larval settlement and (a) crustose coralline algae at Florence Bay (lower turbidity) on downwards-facing surfaces, (b) tubes worms at Picnic Bay (higher turbidity) on downward-facing surfaces, and (c) empty space at Picnic Bay (higher turbidity) on upwards-facing surfaces.

### Fig. S3 Community composition on smothered plugs

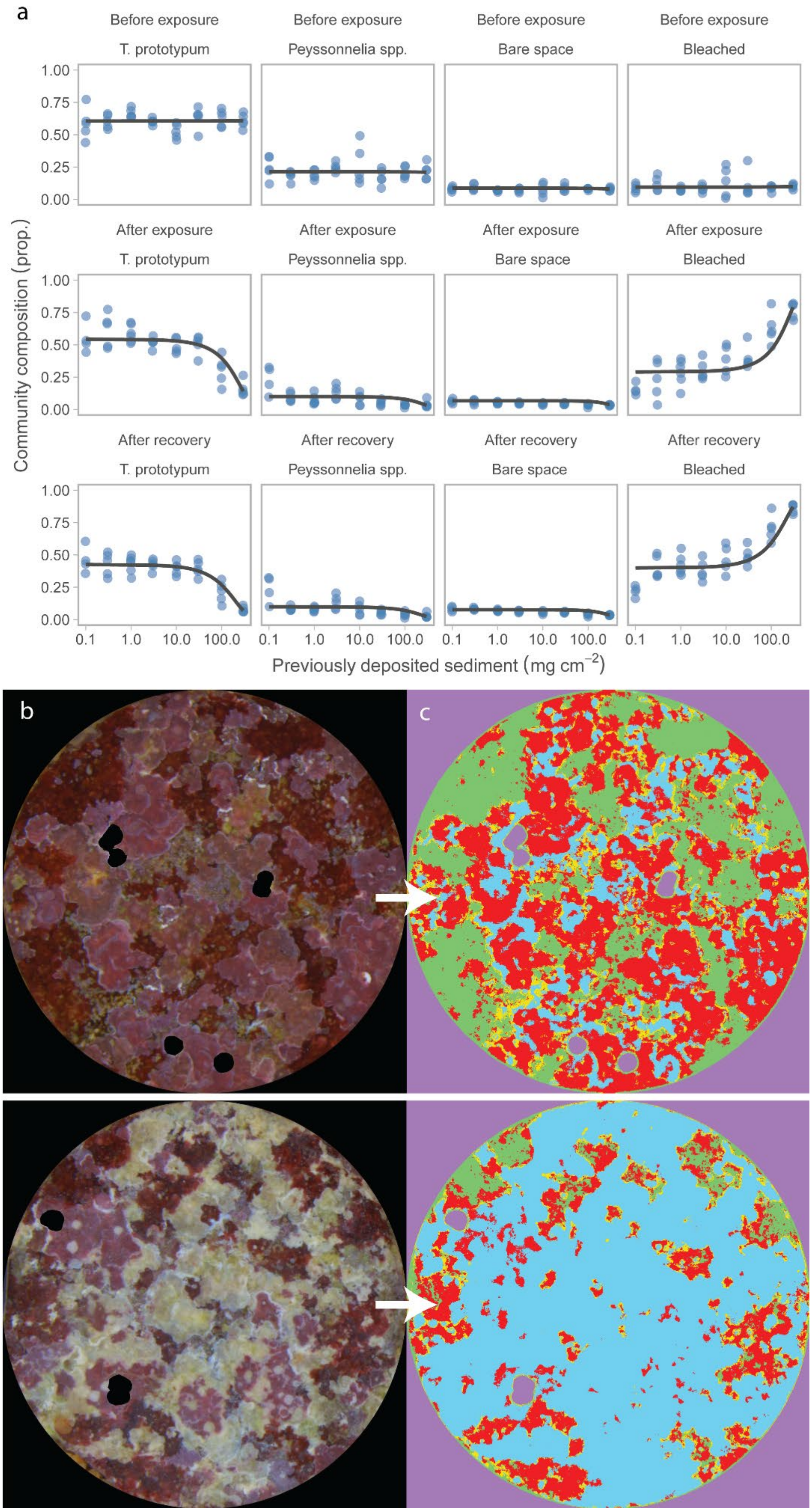

Fig. S3. The classification of algae, bleaching and bare space on settlement plugs before, after and following recovery of exposure and removal of deposited sediments. (a) Community composition of the plugs after each exposure. (b) A representative control plug (no sediment) and treatment plug (100 mg $cm^{-2}$) after sediment removal, and their corresponding segmented images.

### Table S1. Settlement orientation

Table S1. Settlement orientations between field and laboratory experiments, and possible confounding factors that could complicate interpretations. This review is not extensive and only is used to highlight the range of approaches and responses in settlement assays.

| Reference | Orientation preference | Sediment likely | Field/lab | Dimensions | Randomly assigned substrates prior to settlement | Condition duration | Substrate community | Location | Spp./Genera |
|---|---|---|---|---|---|---|---|---|---|
| Babcock, R., & Mundy, C. (1996). Coral recruitment: Consequences of settlement choice for early growth and survivorship in two scleractinians. *Journal of Experimental Marine Biology and Ecology*, *206*(1–2), 179–201. | *Platygyra sinensis* = Vertical at all depths (field). Vertical at both light intensities (lab).<br>*Oxypora lacera* = Undersides at shallow (0 m), vertical at mid (1.5 m) and deep (4.5 m) sites (field). Undersides in higher light, upwards and vertical at lower light (lab). | Unlikely as enclosed in plastic bowls and 200 µm mesh covered openings or in aquaria. | Field enclosures, and laboratory. | 8 cm (upwards) x 8 cm (downward) x 2 cm (vertical) | Non-random (field). Unreported (lab). | 2 months in aquaria and 2 weeks in field | Unreported | Magnetic Island, GBR | *Platygyra sinensis* and *Oxypora lacera* |
| Bak R, Engel M (1979) Distribution, abundance and survival of juvenile hermatypic corals (Scleractinia) and the importance of life history strategies in the parent coral community. Mar Biol 54:341-352 | Vertical in shallow water (3–17 m) and horizontal in deep water (17–37 m) | Possible in all locations | Field | Surveyed reef substrate using 1 x 1 m quadrats | Non-random | Not described | Reef bottom | Curaçao and Bonaire (southern Caribbean) | *Agaricia agaricites* |
| Davidson J, Thompson A, Logan M, Schaffelke B. (2019) High spatio-temporal variability in Acroporidae settlement to inshore reefs of the Great Barrier Reef. PLOS ONE. 2019;14(1):e0209771. | Vertical surfaces, especially once standardised for surface area | Possible in all locations | Field | 11.5 (upwards x 11.5 (downwards) x 1.1 cm (vertical). | Non-random | Min 10 days | Predominantly Biofilm | GBR | *Acropora spp.* |

| | | | | | | | | | |
|---|---|---|---|---|---|---|---|---|---|
| Ricardo GF, Jones RJ, Nordborg M, Negri AP.(2017) Settlement patterns of the coral *Acropora millepora* on sediment-laden surfaces. Sci Total Environ. 2017;609:277-88. | Upwards facing-surfaces. Changed to underside as sediment increased | Used as treatments | Laboratory | 4 x 2 cm diameter each side (upwards, upwards-side, vertical, downwards side, downwards) | Random | 2–3 months | Predominantly *Titanoderma prototypum* CCA | Great Barrier Reef | *Acropora millepora* |
| Erwin P, Song B, Szmant A, (2008) Settlement behaviour of *Acropora palmata* planulae: Effects of biofilm age and crustose coralline algal cover. Proceedings of the 11th international coral reef symposium Ft Lauderdale, Florida; 2008. | Undersides | No but possible during conditioning | Lab (tiles conditioned in field) | Not reported | Non-random | Conditioned in field for 2, 4, 6, 8 and 9 weeks | CCA (0–12%) and biofilm | Caribbean | *Acropora palmata* |
| Lal R, Kininmonth S, N'Yeurt AD, Riley RH, Rico C. (2018) The effects of a stressed inshore urban reef on coral recruitment in Suva Harbour, Fiji. Ecology and Evolution. 8(23):11842-56. | Downward-facing surfaces (only these had grooves) | Yes during conditioning and settlement | Field | 12 (upwards) x 12 (downwards) x 1 cm (vertical) | Non-random | 1 month in field | CCA – exact coverage not known | Fiji | *Acroporidae, Pocilloporidae, Poritidae, and Lobophyllidae* |
| Nozawa, Y., Tanaka, K., & Reimer, J. D. (2011). Reconsideration of the surface structure of settlement plates used in coral recruitment studies. *Zoological Studies*, *50*(1), 53–60. | Upwards-facing surfaces on plates with micro-crevices.<br><br>Downward-facing surfaces on plates without micro-crevices | Possible but settlement plates fixed at steep angle to reduce influence of sediment | Field | 10 × 10 × 0.5 cm flat and grooved plates | Non-random | 1 month | Predominantly CCA | Japan | *Pocilloporids* |
| Price N. (2010) Habitat selection, facilitation, and biotic settlement cues affect distribution and performance of coral recruits in French Polynesia. Oecologia. 2010;163(3):747-58. | Downward-facing surfaces (especially in cryptic underside habitat) | Yes | Field | 15 (upwards) x 15 (downwards) x 15 cm (vertical) | Non-random | 1 month | CCA | French Polynesia | *Acropora, Porites, Montastrea, Favia,* or *Siderastrea* |
| Raimondi PT and Morse ANC (2000) The consequences of complex larval behavior in a coral. Ecology 81:3193-3211 | Downward-facing surfaces | Unlikely as enclosed in chamber | Field | ~ 3 cm diameter upwards, vertical, downwards | Random | NA (artificial cue) | *Hydrolithon sp.* resin | Bonaire (southern Caribbean) | *Agaricia humilis* |
| Szmant AM, Miller MW (2006) Settlement preferences and post-settlement mortality of laboratory cultured and settled larvae of the Caribbean hermatypic corals *Montastraea faveolata* and *Acropora* | Species differences<br><br>*M. faveolata* preferred downward-facing conditioned surfaces, but | Unlikely | Lab | Upwards and downwards, a range of substrate types and sizes. | Inverted | Not reported | For *M. faveolata* – coral rubble, field conditioned clay tiles and field conditioned limestone plates. | Florida keys | *Montastraea faveolata* and *Acropora palmata* |

| | | | | | | | | | |
|---|---|---|---|---|---|---|---|---|---|
| *palmata* in the Florida Keys, USA. Proc 10th Int Coral Reef Sym 1:43-49 | preferred upwards if the plate was inverted.<br><br>*A. palmata* had no preference or preferred upwards-facing surfaces even if they were inverted. | | | | | | For *A. palmata* – field conditioned limestone plates<br><br>All with predominant CCA cover | | |
| Vermeij M. (2006) Early life-history dynamics of Caribbean coral species on artificial substratum: the importance of competition, growth and variation in life-history strategy. Coral Reefs. 2006;25(1):59-71. | 11 out of 17 species preferred upwards-facing surfaces<br><br>6 out of 17 species preferred downwards facing surfaces.<br><br>Most recruits found on downwards facing surfaces driven by 4 dominant spp. | Likely. Although the design would allow for some sediment removal. | Field | Egg crate 120 x 60 cm position upwards, downwards and vertically. | Non-random | ~1 year in situ<br><br>Surveyed every year for 6 years | Benthic reef components (including CCA, coral, sponges macroalgae) | Curacao (Caribbean) | 17 scleractinian species |
| Yusuf S, Zamani N, Jompa J, Junior M, editors. Larvae of the coral *Acropora tenuis* (Dana 1846) Settle under controlled light intensity. (2019) IOP Conference Series: Earth and Environmental Science; 2019: IOP Publishing. | Horizontal | No | Laboratory | 10 (upwards facing) x 10 cm (vertical). | Non-random | Not reported | Artificial substrate | GBR | *Acropora tenuis* |
