## Supplementary material for "Impacts of water quality on *Acropora* coral settlement: The relative importance of substrate quality and light": Highlights

- Light intensity and spectral changes overall had limited impact on coral settlement success.
- Monochromatic/artificial light exposures led to changes in settlement at higher wavelengths.
- Substrates conditioned under various water quality exposures had a greater impact on coral settlement.
