## Supplementary figures and images for "Impacts of water quality on *Acropora* coral settlement: The relative importance of substrate quality and light"

### Graphical Abstract

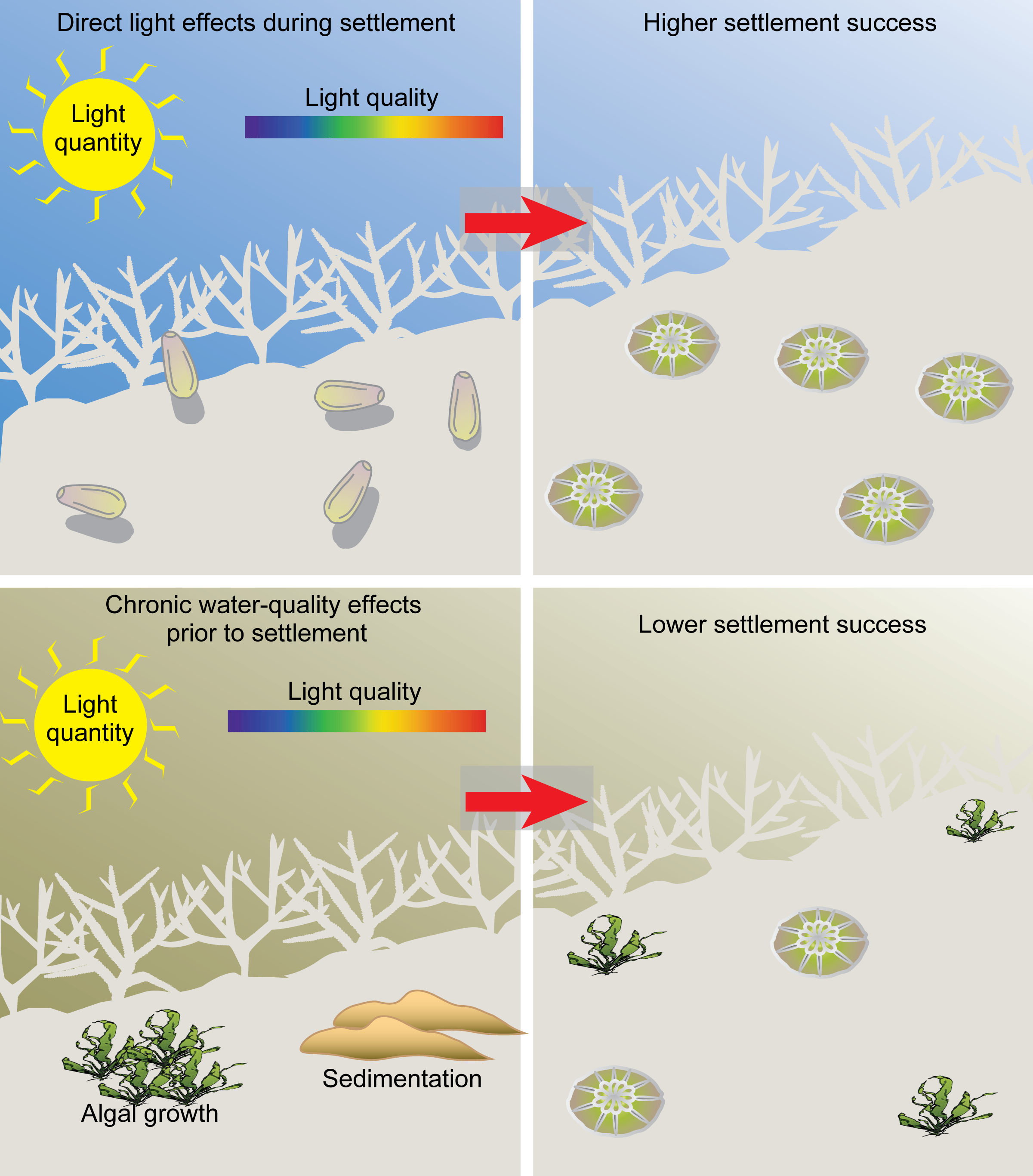
